# The jellyfish genome sheds light on the early evolution of active predation

**DOI:** 10.1101/449082

**Authors:** Hak-Min Kim, Jessica A. Weber, Nayoung Lee, Seung Gu Park, Yun Sung Cho, Youngjune Bhak, Nayun Lee, Yeonsu Jeon, Sungwon Jeon, Victor Luria, Amir Karger, Marc W. Kirschner, Ye Jin Jo, Seonock Woo, Kyoungsoon Shin, Oksung Chung, Jae-Chun Ryu, Hyung-Soon Yim, Jung-Hyun Lee, Jeremy S. Edwards, Andrea Manica, Jong Bhak, Seungshic Yum

## Abstract

**Background:** Unique among cnidarians, jellyfish have remarkable morphological and biochemical innovations that allow them to actively hunt in the water column. One of the first animals to become free-swimming, jellyfish employ pulsed jet propulsion and venomous tentacles to capture prey.

**Results:** To understand these key innovations, we sequenced the genome of the giant Nomura’s jellyfish (*Nemopilema nomurai*), the transcriptomes of its bell and tentacles, and transcriptomes across tissues and developmental stages of the *Sanderia malayensis* jellyfish. Analyses of *Nemopilema* and other cnidarian genomes revealed adaptations associated with swimming, marked by codon bias in muscle contraction and expansion of neurotransmitter genes, along with expanded Myosin type II family and venom domains; possibly contributing to jellyfish mobility and active predation. We also identified gene family expansions of *Wnt* and posterior *Hox* genes, and discovered the important role of retinoic acid signaling in this ancient lineage of metazoans, which together may be related to the unique jellyfish body plan (medusa formation).

**Conclusions:** Taken together, the jellyfish genome and transcriptomes genetically confirm their unique morphological and physiological traits that have combined to make these animals one of the world’s earliest and most successful multi-cellular predators.

## Background

Cnidarians, including jellyfish and their predominantly sessile relatives the coral, sea anemone, and hydra, first appeared in the Precambrian Era and are now key members of aquatic ecosystems worldwide [1]. Between 500 and 700 million years ago, jellyfish developed novel physiological traits that allowed them to become one of the first free-swimming predators. The life cycle of the jellyfish includes a small polypoid, sessile stage which reproduces asexually to form the mobile medusa form that can reproduce both sexually and asexually [2]. The class Scyphozoa, or true jellyfish, are characterized by a predominant medusa life-stage consisting of a bell and venomous tentacles used for hunting and defense [3]. Jellyfish medusae feature a radially symmetric body structure, powered by readily identifiable cell types such as motor neurons and striated muscles that expand and contract to create the most energy-efficient swimming method in the animal kingdom [4, 5]. Over 95% water, jellyfish are osmoconformers that use ion gradients to deliver solutes to cells and tissues where sodium and calcium ions activate the muscle contractions that power their propulsion. Notably, many jellyfish species can survive in habitats with varying levels of salinity and are successful in low-oxygen environments, allowing them to bloom even in dead zones [6]. These innovations have allowed them to colonize aquatic habitats across the globe both in brackish and marine environments, spanning the shallow surface waters to the depths of the seas.

## Results and discussion

### Jellyfish genome assembly and annotation

Here, we present the first *de novo* genome assembly of a jellyfish (*Nemopilema nomurai*). It resulted in a 213 Mb genome comprised of 255 scaffolds and an N50 length of 2.71 Mb, containing only 1.48 % gaps (Additional file 1: Tables S2 and S3). The *Nemopilema* hybrid assembly was created using a combination of short and long read sequencing technologies, consisting of 38.2 Gb Pacific Biosciences (PacBio) single molecule real time sequencing (SMRT) reads, along with 98.6 Gb of Illumina short insert, mate-pair, and TruSeq synthetic long reads (Additional file 1: Figures S3-S5; Tables S4-S7). The resulting assembly shows the longest continuity among cnidarian genomes (Additional file 1: Table S9). We predicted 18,962 protein-coding jellyfish genes by combining *de novo* (using medusa bell and tentacle tissue transcriptomes) and homologous gene prediction methods (Additional file 1: Tables S10 and S11). This process recovered the highest number of single-copy orthologous genes [7] among all published non-bilaterian metazoan genome assemblies to date (Additional file 1: Table S12). A total of 21.07% of the jellyfish genome was found to be made up of transposable elements, compared to those of *Acropora digitifera* (9.45%), *Nematostella vectensis* (33.63%), and *Hydra magnipapillata* (42.87%) (Additional file 1: Table S13).

We compared the *Nemopilema* genome to other cnidarian genomes, all of which are from predominantly sessile taxa, to detect unique Scyphozoa function (active mobility), physical structure (medusa bell), and chemistry (venom). We also performed transcriptome analyses of both *Nemopilema nomurai* and the *Sanderia malayensis* jellyfish across three medusa tissue types and four developmental stages.

### Evolutionary analysis of the jellyfish

To identify jellyfish-specific evolutionary traits, we examined gene family expansions and contractions across one unicellular holozoan and eleven metazoans using 15,255 orthologous gene families (see Additional file 1: Section 4.1). Of these, 7,737 were found in *Nemopilema* and 4,156 were shared by all four available cnidarian genomes (*Nemopilema nomurai*, *Hydra magnipapillata*[8], *Acropora digitifera*[9], and *Nematostella vectensis*[10]; Fig. 1a). A phylogeny constructed using these orthologs revealed a monophyletic cnidarian clade that diverged from the metazoan stem prior to the evolution of the bilaterians (Fig. 1b; Additional file 1: Figure S7). To determine how many genes appeared in every evolutionary era in the genome of Nomura’s jellyfish, we also evaluated the evolutionary age of the protein-coding genes. Grouping jellyfish genes into 3 broad evolutionary eras, we observed that while the majority (80%) of genes are ancient (older than 741 Mya), a few (~3%) are of an intermediate age (741 - 239 Mya) and some (17%) are young (239 Mya to present; Fig. 1c; Additional file 1: Figure S10). Interestingly, normalizing the number of genes by the age and length of evolutionary era suggests that gene turnover is highest near the present time. In total, the *Nemopilema* genome contained 67 expanded and 80 contracted gene families compared to the common ancestor of *Nemopilema* and *Hydra* (Fig. 1b; see Additional file 1: Section 4.2). Gene Ontology (GO) terms related to sensory perception were under-represented in the Cnidaria lineage compared to Bilateria, accurately reflecting cnidarian’s less complex sensory system (Additional file 1: Tables S14 and S15). However, neurotransmitter transport function (GO:0005326, *P* = 1.66E-16) was significantly enriched in *Nemopilema* compared to other cnidarians (Additional file 1: Tables S16 and S17), likely due to the balance and visual structures, such as the statocyst and ocelli, that are more elaborate in the mobile medusa than in sessile polyps [11].

**Fig. 1.**
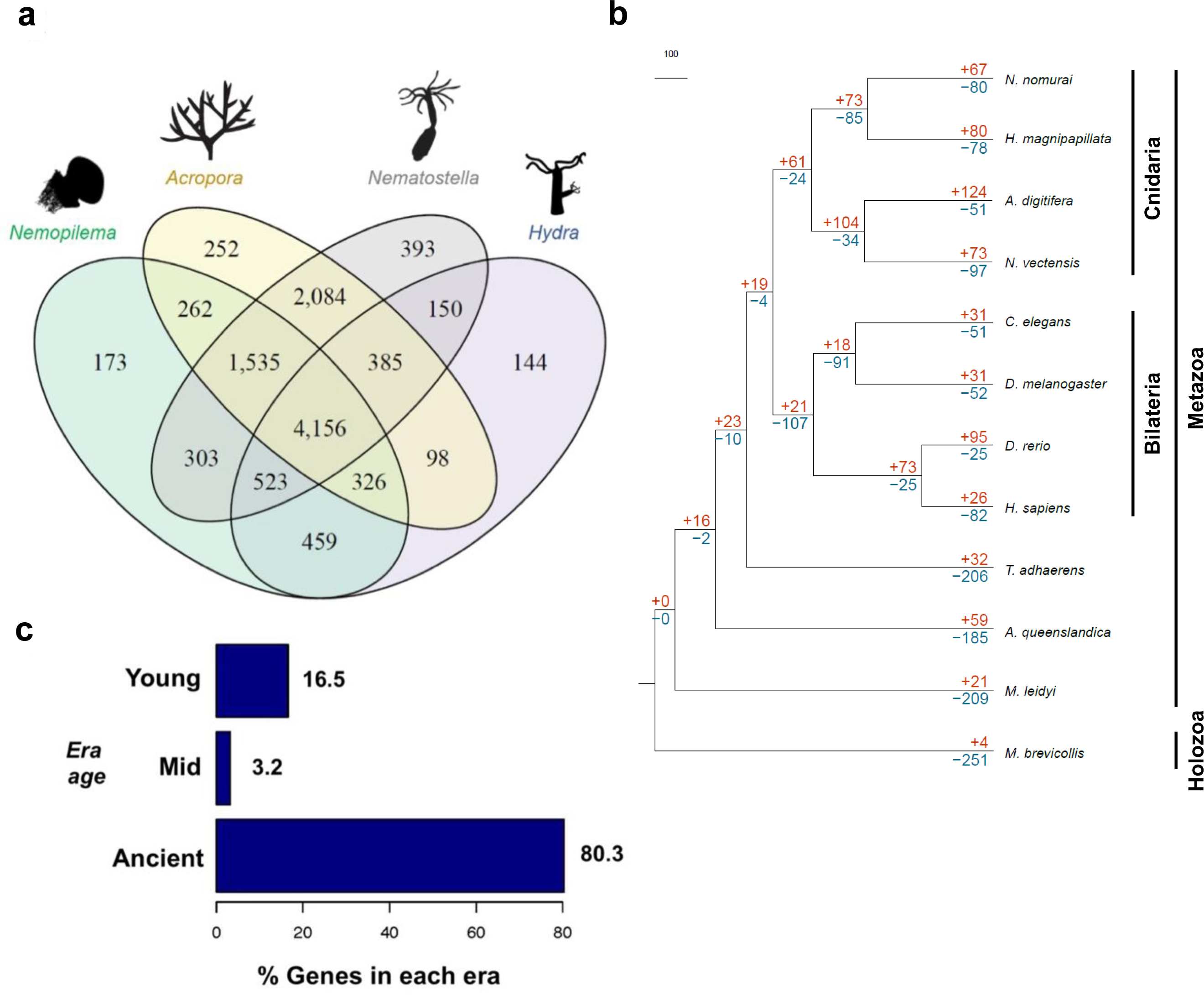
Gene family relationships of cnidarian and metazoan species. **a** Venn diagram of the number of unique and shared gene families among four cnidarian genomes. **b** Gene family expansions and contractions in the *Nemopilema* genome. Numbers designate the number of gene families that have expanded (red, +) and contracted (blue, -) after the split from the common ancestor. **c** The proportion of *Nemopilema* genes in each evolutionary era. While most *Nemopilema* genes (~80%) are ancient (~1,877 Mya), a few (~3%) are of intermediate age (~659 Mya) and a significant fraction (~17%) are relatively young (~147 Mya).

### Genomic context and muscle associated genes

Jellyfish have two primary muscle types: the epitheliomuscular cells, which are the predominant muscle cells found in sessile cnidarians; and the striated muscle cells located in the medusa bell that are essential for swimming. To understand the evolution of active-swimming in jellyfish, we examined their codon bias compared to other metazoans by calculating the guanine and cytosine content at the third codon position (GC3) [12, 13] (Additional file 1: Figure S13). It has been suggested that genes with high level of GC3 are more adaptable to external stresses (e.g., environmental changes) [14]. Among the high-scoring top 100 GC3 biased genes, the regulation of muscle contraction and neuropeptide signaling pathways GO terms were specific to *Nemopilema* (Additional file 4). Calcium plays a key role in the striated muscle contraction in jellyfish, and the calcium signaling pathway (GO:0004020, *P* = 5.60E-10) showed a high level of GC3 biases specific to *Nemopilema. Nemopilema* top 500 GC3 genes were enriched in GO terms associated with homeostasis (e.g. cellular chemical homeostasis and sodium ion transport), which we speculate is essential for the activation of muscle contractions that power the jellyfish’s mobile predation (see Additional file 1: Section 5.1).

Since cnidarians have been reported to lack titin and troponin complexes, which are critical components of bilaterian striated muscles, it has been suggested that the two clades independently evolved striated muscles [15]. A survey of genes that encode muscle structural and regulatory proteins in cnidarians showed a conserved eumetazoan core actin-myosin contractile machinery shared with bilaterians (Additional file 1: Table S23). However, like other cnidarians, *Nemopilema* lacks titin and troponin complexes, which are key components of bilaterian striated muscles. Also, γ-syntrophin, a component of the dystroglycan complex, was absent in both *Nemopilema* and *Hydra*. However, *Nemopilema* do possess α/β-Dystrobrevin and α/∊-Sarcoglycan dystroglycan-associated costamere proteins, indicating that several components of the dystroglycan complex were lost after the Scyphozoa-Hydrozoa split. It was suggested that *Hydra* undergone secondary simplifications relative to *Nematostella*, which has a greater degree of muscle-cell-type specialization [8]. Compared to *Hydra* and *Nematostella*, *Nemopilema* shows intermediate complexity of muscle structural and regulatory proteins between *Hydra* and *Nematostella*.

### Medusa bell and tentacle transcriptome profiling

Jellyfish medusa bell and tentacles are morphologically distinct and perform discrete physiological functions [16, 17]. We generated bell and tentacle transcriptomes from *Nemopilema* and the smaller *Sanderia malayensis,* which can be grown in the laboratory, to assess developmental regulation (Additional file 1: Table S20). Enrichment tests of highly expressed genes showed that muscle-associated functional categories (e.g. muscle myosin complex and muscle tissue morphogenesis) were enriched in the bell (Fig. 2a; see Additional file 5). Myosins comprise a superfamily of motor proteins and play a critical role in muscle contraction and are involved in a wide range of motility processes in Eukaryotes. Critically, the Myosin II family proteins, found in cells of both striated muscle tissue and smooth muscle tissue, are responsible for producing contraction in muscle cells [18]. Cnidarians possess both epitheliomuscular cells and striated muscle cells. Striated muscle is a critical component of the subumbrella of the medusa bell, where its fast contractions power the unique propulsion-based swimming of the jellyfish. We found that type II Myosin heavy chain (MYH) and Myosin light chain (MYL) gene families were highly expressed in the bell, and are closely associated with striated and smooth muscle cells [15]. Interestingly, *Nemopilema* also showed the largest copy numbers of MYH and MYL genes among non-bilaterian metazoans (Fig. 2c; see Additional file 1: Section 5.3), and six of the seven MYH genes and 12 out of 21 MYL genes showed higher expression in the bell than the tentacles with very high ~8.8 and ~17-fold increase on average, respectively (Fig. 2d). These results suggest that the combinations of copy number expansion of type II Myosin gene families and high expression of muscle associated genes confirmed that muscles in medusa bell are an important determinant of jellyfish motility.

**Fig. 2.**
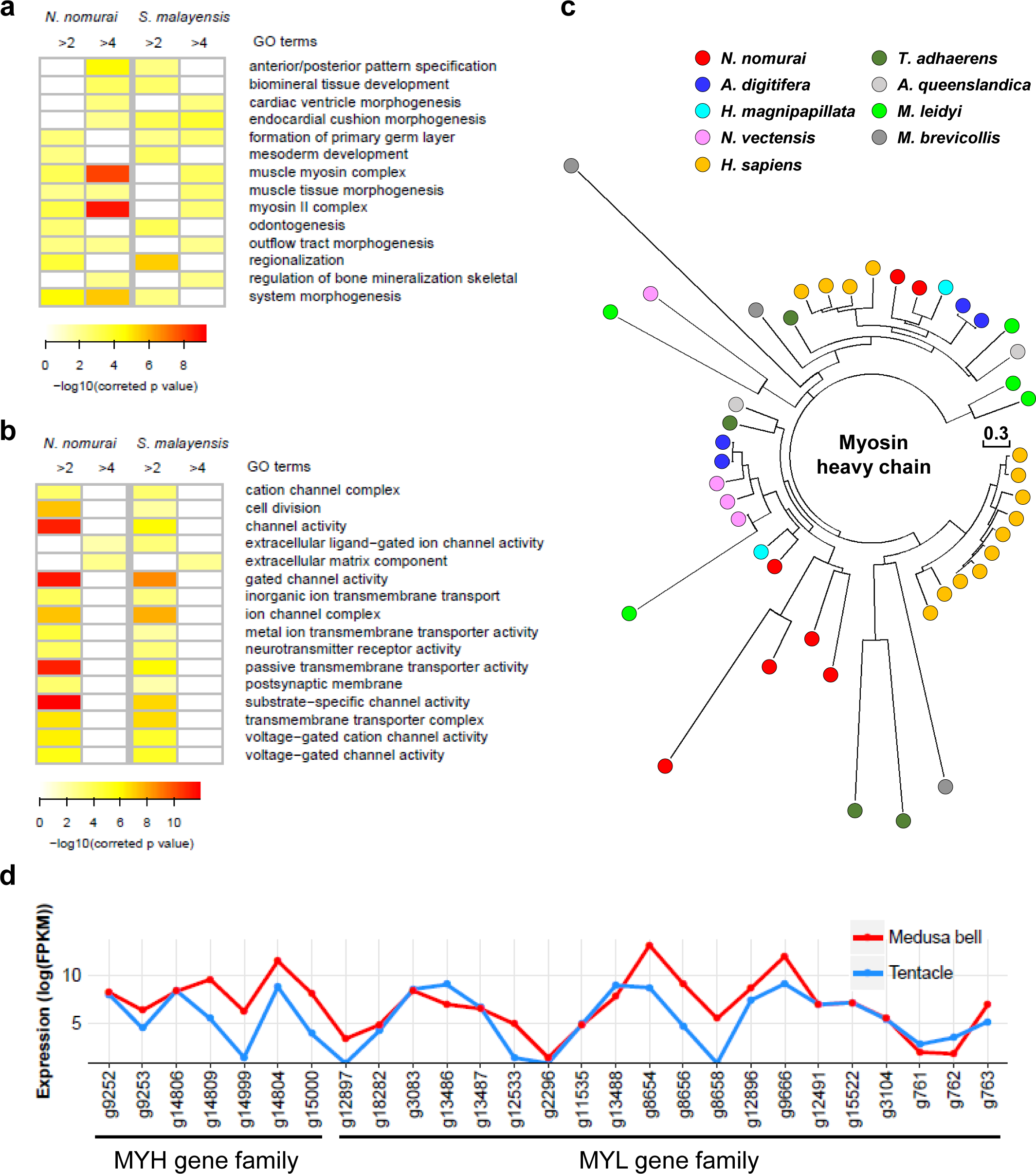
Gene expression patterns of medusa bell and tentacle tissues and expansion of myosin heavy chain genes in jellyfish. **a** *P*-value heatmap of enriched GO categories using highly expressed genes in medusa bell tissue. Greater than 2-fold and 4-fold higher expression in medusa bell than tentacles are shown in each column. Only shared GO categories between *N. nomurai* and *S. malayensis* are shown. **b** *P*-value heatmap of enriched GO categories using highly expressed genes in tentacle tissue. **c** Unrooted Le-Gascuel model tree of myosin heavy chain genes using BLAST best hit method. **d** Expression pattern of MYH and MYL genes in *Nemopilema*. Genes that are not expressed in both tentacles and medusa bell were excluded.

Conversely, gene expression analyses of the tentacles revealed high RNA expression levels of neurotransmitter associated functional categories (ion channel complex, postsynapse, and neurotransmitter receptor activity; Fig. 2b); consistent with the anatomy of jellyfish tentacles, which contain the sensory cells and a loose plexus of the neuronal subpopulation at the base of the ectoderm [19].

### Body patterning in the jellyfish

There has been much debate surrounding the early evolution of body patterning in the metazoan common ancestor, particularly concerning the origin and expansion of Hox and *Wnt* gene families [20-22]. In total, 83 homeodomains were found in *Nemopilema*, while 41, 120, and 148 of homeodomains were found from *Hydra*, *Acropora*, and *Nematostella*, respectively (Additional file 1: Table S24). Five of the eight Hox genes in *Nemopilema* are of the posterior type that are associated with aboral axis development [22] and clustered with *Nematostella’s* posterior Hox genes, *HOXE* and *HOXF*(Additional file 1: Figures S18-S20). Though absent in *Hydra* and *Acropora,* synteny analyses of ParaHox genes in *Nemopilema* show that the *XLOX/CDX* gene is located immediately downstream of *GSX* in the same tandem orientation as those in *Nematostella,* suggesting that *XLOX/CDX* was present in the cnidarian common ancestor and subsequently lost in some lineages (Additional file 1: Figure S21). Hox related genes, *EVX* and *EMX,* are also present in *Nemopilema*, although they are absent in *Hydra*. Given the large amount of ancestral diversity in the *Wnt* genes, it has been proposed that Wnt signaling controlled body plan development in the early metazoans [23]. *Nemopilema* possesses 13 *Wnt* orthologs representing 10 *Wnt* subfamilies (Additional file 1: Figure S22; Table S25).*Wnt9* is absent from all cnidarians, likely representing losses in the cnidarian common ancestor. Cnidarians have undergone dynamic lineage specific *Wnt* subfamily duplications, such as *Wnt8* (*Nematostella* and *Acropora*), *Wnt10* (*Hydra*), and *Wnt11*, and *Wnt16* (*Nemopilema*). It has been proposed that a common cluster of *Wnt* genes (*Wnt1*–*Wnt6*–*Wnt10*) existed in the last common ancestor of arthropods and deuterostomes [24]. Our analyses of cnidarian and bilaterian genomes revealed that *Acropora* also possess this cluster, while *Nemopilema* and *Hydra* are missing *Wnt6*, suggesting loss of the *Wnt6* gene in the Medusozoa common ancestor (Additional file 1: Figure S23). Taken together, the jellyfish have comparable number of Hox and *Wnt* genes to other cnidarians, but the dynamic repertoire of these gene families suggests that cnidarians have evolved independently to adapt their physiological characteristics and life cycle.

### Polyp to medusa transition in jellyfish

The polyp-to-medusa transition is prominent in jellyfish compared to the other sessile cnidarians. To understand the genetic basis of the medusa structure formation in the jellyfish, we compared transcriptional regulation between cnidarians and across jellyfish developmental stages (see Additional file 1: Sections 7.1 and 7.2). We assembled the *Sanderia* transcripts using six pooled samples of transcriptomes (Additional file 1: Table S26). The assembled transcripts had a total length of 61 Mb and resulted in 58,290 transcript isoforms and 43,541 unique transcripts, with a N50 of 2,325 bp. On average, 87% of the RNA reads were aligned to into the assembled transcripts (Additional file 1: Table S27), indicating that the transcript assembly represented the majority of sequenced reads. Furthermore, the composition of the protein domains contained in the top 20 ranks was quite similar between *Nemopilema* and *Sanderia* (Additional file 1: Table S28). To obtain differentially expressed genes for each stage, we compared each stage with the previous or next stage in the life cycle of the jellyfish. The polyp stage, which represents a sessile stage in the jellyfish life cycle, showed enriched terms related to ion channel activity and energy metabolism (regulation of metabolic process, and amino sugar metabolic process; Additional file 1: Table S29). Active feeding in the polyp stimulates asexual proliferation either into more polyps or metamorphosis to strobila [25]. Since anthozoans do not form a medusa, the strobila asexual reproductive stage is an important stage in which to study the metamorphosis from polyp to medusa. In this stage, GO terms related to amide biosynthetic and metabolic process were highly expressed compared to the polyp stage (Additional file 1: Table S30). It has been reported that RF-amide and LW-amide neuropeptides were associated with metamorphosis in cnidarians [26-28]. However, we could not confirm this finding in our strobila and ephyra stage comparisons. In our system, the gene expression patterns of the two stages are quite similar. In the ephyra, the released mobile stage, GO terms involving amide biosynthetic and metabolic process were also highly expressed compared to the merged medusa stage (Additional file 1: Table S31). In the medusa, extracellular matrix, metallopeptidase activity, and immune system process terms were enriched (Additional file 1: Table S32), consistent with the physiology of their bell, tentacles, and oral arm tissue types.

Polyp-to-medusa metamorphosis was previously shown to be strongly associated with *CL390* and *retinoid X receptor*(*RXR*) genes in the *Aurelia aurita* jellyfish [29]. Interestingly, *CL390* was not found in *Nemopilema* or other published cnidarians, suggesting that it may be an *Aurelia*-specific strobilation inducer gene. However, we confirm that *RXR* is present in *Nemopilema*, and absent from cnidarians without a prominent medusa stage (Additional file 1: Figure S24). Retinoic acid (RA) signaling plays a central role during vertebrate growth and development [30], where it regulates transcription by interacting with the RA receptor (RAR) bound to RA response elements (RAREs) of nearby target genes [31]. Of the genes in the RA signaling pathway, *Nemopilema* possess ADH and RALDH enzymes that metabolize retinol to RA, and *RXR* and RAREs to activate transcription of the target gene (Fig. 3a). We discovered 1,630 *Nemopilema* RAREs regions with an average distance of 13 Kbp to the nearest gene (Fig. 3b; Additional file 1: Tables S33 and S34). Interestingly, four posterior Hox genes of *Nemopilema* were located within ±10 Kbp from RAREs, which is unique among the non-bilaterian metazoans (Fig. 3c). Together these findings suggest that retinoic acid signaling was present in early metazoans for regulating target genes with *RXR* and RAREs, and that *RXR* and RAREs may play a critical role for polyp-to-medusa metamorphosis [29]

**Fig. 3.**
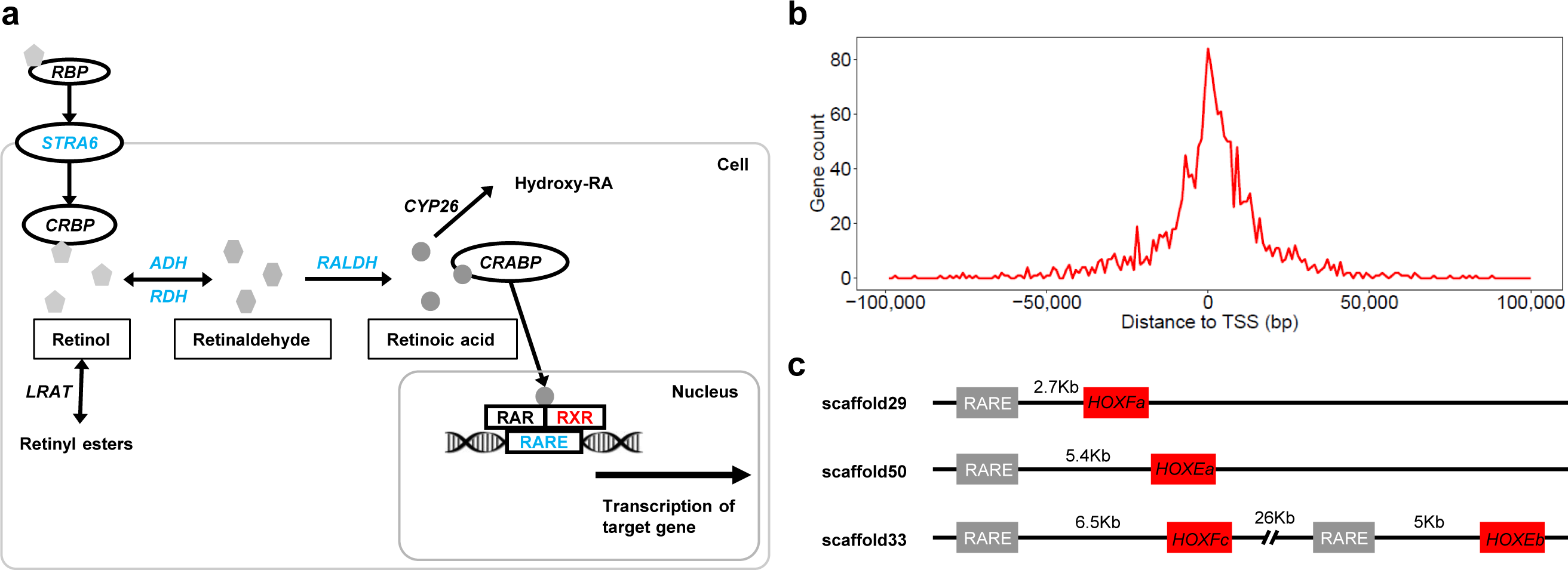
Retinoic acid signaling pathway and RAREs in *Nemopilema*. **a** Schematic of the retinoic acid signaling pathway in humans. Blue denotes presence of the gene and/or element in Cnidaria. Red denotes presence only in *Nemopilema* among the published cnidarians. **b** The distribution of distances between the RAREs and the nearest gene. The distance was calculated by identifying its proximity to transcription start site (TSS) of the genes. The gene count was calculated for each non-overlapping 1 Kb bin across a range of -100 Kb to 100 Kb. **c** The RAREs located nearby posterior Hox genes in *Nemopilema*.

### Identification of toxin related domains in jellyfish

Jellyfish produce complex mixtures of proteinaceous venoms for active prey capture and defense [32]. We identified abundant toxin domains in *Nemopilema* when compared to the non-bilaterian metazoan gene sets in the Tox-Prot database [33]. In total, 69 out of 136 toxin domains aligned to non-bilaterian metazoans; of these 69 toxin domains, 53 were found in *Nemopilema* (Additional file 1: Table S35). Expectedly, the *Nemopilema* genome contains the largest number of venom or toxin associated domains of the included non-bilaterian metazoans. These domains inhibitor domain (PF07648), phospholipase A_2_ (PF05826), and ShK domain-like (PF01549) domains (Fig. 4). Compared to the common ancestor of *Nemopilema* and *Hydra*, *Nemopilema* include Reprolysin (M12B) family zinc metalloprotease (PF01421), Kazal-type serine protease showed expanded gene families associated with metallopeptidase activities (GO:0008237, *P* = 1.99E-16). In particular, Reprolysin (M12B) family zinc metalloproteases are enzymes that cleave peptides and comprise most snake venom endopeptidases [34]. Furthermore, it has been reported that serine protease inhibitor and ShK domains were abundantly found in the transcriptomes of both the cannonball jellyfish (*Stomolophus meleagris*), and the box jellyfish (*Chironex fleckeri*)[35, 36], and phospholipase A_2_ is well-characterized toxin-related enzyme, which is critical to the production of venom components, found in the class Scyphozoa [37].

**Fig. 4.**
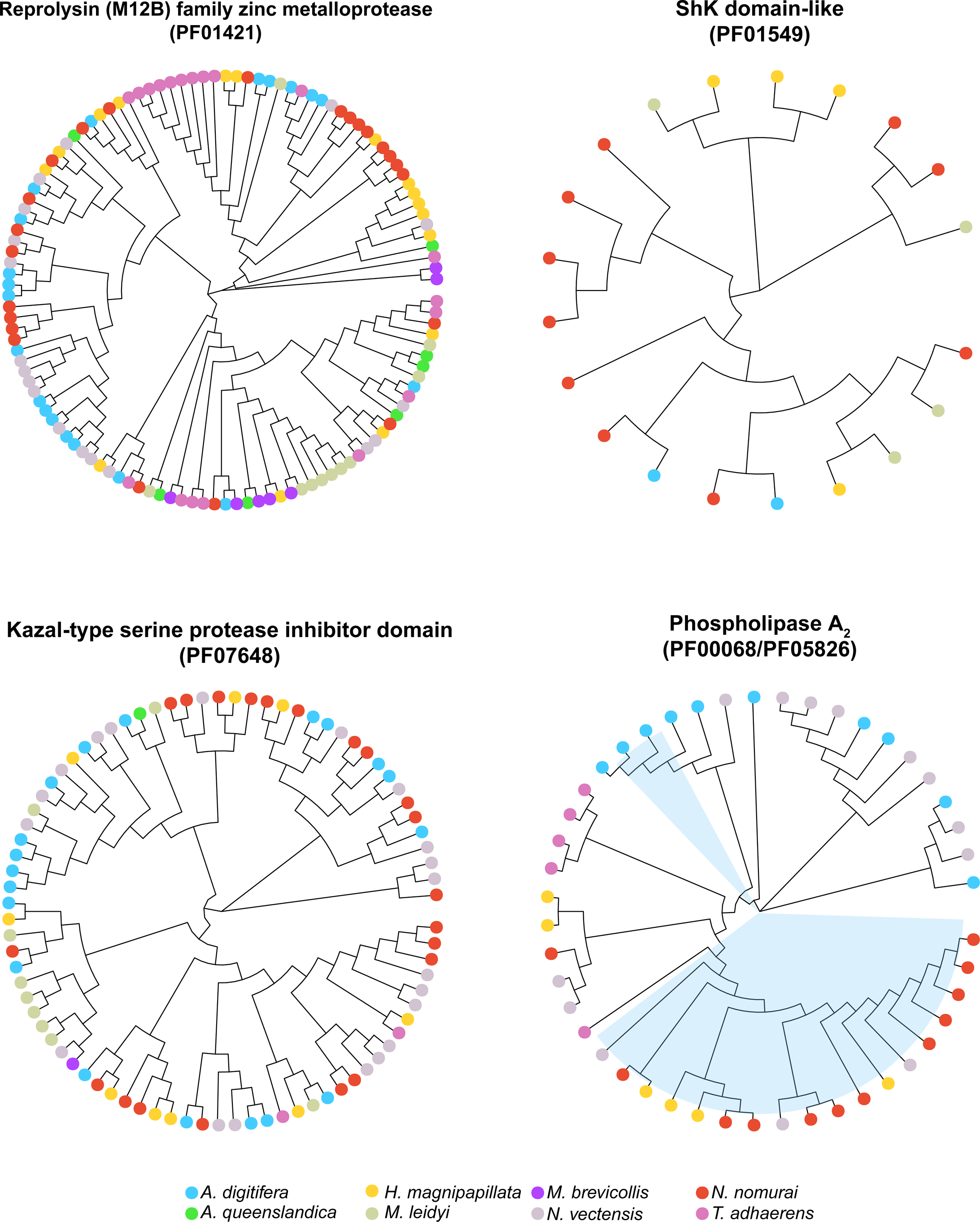
Phylogenetic analysis of venom related domains in non-bilaterian metazoans. Five venom domains (PF01421, PF01549, PF07648, PF00068, and PF05826) are represented in four circular dendrograms. Two phospholipase A_2_ domains (PF00068 and PF05826) were merged into one circular dendrogram (bottom right) and shadings on branches and nodes (sky-blue) in phospholipase A_2_ denote the PF05826 domain.

## Conclusions

A unique branch on the tree of life, jellyfish have evolved remarkable morphological and biochemical innovations that allow them to actively hunt using pulsed jet propulsion and venomous tentacles. While the expansion and contraction of distinct families reflect the adaptation to salinity and predation and the convergent evolution of muscle elements, the *Nemopilema* genome strikes a balance between the conservation of many ancient genes and an innovative potential reflected in significant number of new genes that appeared since *Rhizostomeae* emerged. The *Nemopilema nomurai* genome has provided clues to the genetic basis of the innovative structure, function, and chemistry that have allowed this distinctive early group of predators to colonize the waters of the globe.

## Methods

### Sample preparation

A medusa *Nemopilema nomurai* was collected at the Tongyeong Marine Science Station, KIOST (34.7699 N, 128.3828 E) on Sep. 12, 2013. The *Sanderia malayensis* samples were obtained from Aqua Planet Jeju Hanwha (Seogwipo, Korea) for transcriptome analyses of developmental stages since *Nemopilema* cannot be easily grown in the laboratory. The DNA and RNA preparation of *Nemopilema* and *Sanderia* are described in the Additional file 1: Section 1.1. Species identification of *Nemopilema* was confirmed by comparing the *MT-COI* gene of five species of jellyfish. We aligned *Nemopilema* Illumina short reads (~400 bp insert-size) to the *MT-COI* gene of *Chrysaora quinquecirrha* (NC_020459.1), *Cassiopea frondosa* (NC_016466.1), *Craspedacusta sowerbyi* (NC_018537.1), and *Aurelia aurita* (NC_008446.1) jellyfish with BWA-MEM aligner [38]. Consensus sequences for each jellyfish were generated using SAMtools [39]. The consensus sequence from *C. sowerbyi* was excluded due to low coverage. We conducted multiple sequence alignment using MUSCLE [40] and ran the MEGA v7 [41] neighbor joining phylogenetic tree (gamma distribution) with 1,000 bootstrap replicates. Mitochondrial DNA phylogenetic analyses confirmed the identification of the *Nemopilema* sample as *Nemopilema nomurai*.

### Genome sequencing and scaffold assembly

For the *de novo* assembly of *Nemopilema*, PacBio SMRT and five Illumina DNA libraries with various insert sizes (400bp, 5 Kb, 10 Kb, 15 Kb, and 20 Kb) were constructed according manufacturers’ protocols. The Illumina libraries were sequenced using a HiSeq2500 with read length of 100 bp (400 bp, 15 Kb, and 20 Kb) and a HiSeq2000 with read length of 101 bp (5 Kb and 10 Kb). Quality filtered PacBio subreads were assembled into distinct contigs using the FALCON assembler [42] with various read length cutoffs. To extend contigs to scaffolds, we aligned the Illumina long mate-pair libraries (5 Kb, 10 Kb, 15 Kb, and 20 Kb) to contig sets and extended the contigs using SSPACE [43]. Gaps generated by SSPACE were filled by aligning the Illumina short-insert paired-end sequences using GapCloser [44]. We also generated TSLRs using an Illumina HiSeq2000, which were aligned to scaffolds to correct erroneous sequences and to close gaps using an in-house script. Detailed genome sequencing and assembly process are provided in Additional file 1: Section 2.2.

### Genome annotation

The jellyfish genome was annotated for protein-coding genes and repetitive elements. We predicted protein-coding genes using a two-step process, with both homology and evidence-based prediction. Protein sequences of the sea anemone, hydra, sponge, human, mouse, and fruit fly from the NCBI database, and Cnidaria protein sequences from the NCBI Entrez protein database were used for homology-based gene prediction. Two tissue transcriptomes from *Nemopilema* were used for evidence-based gene prediction via AUGUSTUS [45]. Final *Nemopilema* protein-coding genes were determined using AUGUSTUS with exon (from homology-based gene prediction) and intron (from evidence-based gene prediction) hints. Repetitive elements were also predicted using Tandem Repeats Finder [46] and RepeatMasker [47]. Details of the annotation process are provided in Additional file 1: Sections 3.1 and 3.2.

### Gene age estimation

Phylostratigraphy employs BLASTP-scored sequence similarity to estimate the minimal age of every protein-coding gene. The protein sequence is used to query the NCBI non-redundant database and detect the most distant species in which a sufficiently similar sequence is present, and inferring that the gene is at least as old as the age of the common ancestor [48]. For every species, we use the NCBI taxonomy. The timing of most divergence events is estimated using TimeTree [49] and the Encyclopedia of Life [50]. To facilitate detection of sequence similarity, we use the e-value threshold of 10^-3^. We evaluate the age of all proteins whose length is equal or greater than 40 amino acids. We count the number of genes in each phylostratum, from most ancient (PS 1) to newest (PS 11). To see broad evolutionary patterns, we aggregate the counts from several phylostrata into 3 broad evolutionary eras: ancient (PS 1-5, cellular organisms to Eumetazoa, 4,204 Mya - 741 Mya), middle (PS 6-7, Cnidaria to Scyphozoa, 741 Mya - 239 Mya) and young (PS 8-11, Rhizostomeae to *Nemopilema nomurai*, 239 Mya to present).

### Comparative evolutionary analyses

Orthologous gene clusters were constructed to examine the conservation of gene repertoires among the genomes of the *Nemopilema nomurai*, *Hydra magnipapillata*, *Acropora digitifera*, *Nematostella vectensis*, *Caenorhabditis elegans*, *Danio rerio*, *Drosophila melanogaster*, *Homo sapiens*, *Trichoplax adhaerens*, *Amphimedon queenslandica*, *Mnemiopsis leidyi*, and *Monosiga brevicollis* using OrthoMCL [51]. To infer a phylogeny and divergence times, we used RAxML [52] and MCMCtree [53], respectively. A gene family expansion and contraction analysis was conducted using the Café program [54]. Domain regions were predicted by InterProScan [55] with domain databases. Details of the comparative analysis are provided in Additional file 1: Sections 4.1-4.4.

### Transcriptome sequencing and expression profiling

Illumina RNA libraries from *Nemopilema nomurai* and *Sanderia malayensis* were sequenced using a HiSeq2500 with 100 bp read lengths. Since there is not a reference genome for *S. malayensis*, we *de novo* assembled a pooled six RNA-seq read set using the Trinity assembler [56]. Quality filtered RNA reads from *Nemopilema* and *Sanderia* were aligned to the *Nemopilema* genome assembly and the assembled transcripts, respectively, using the TopHat [57] program. Expression values were calculated by the Fragments Per Kilobase Of Exon Per Million Fragments Mapped (FPKM) method using Cufflinks [57], and differentially expressed genes were identified by DEGseq [58]. Details of the transcriptome analysis are presented in Additional file 1: Sections 5.2 and 7.1.

### Hox and ParaHox analyses

We examined the homeodomain regions in *Nemopilema* using the InterProScan program. Hox and ParaHox genes were identified in *Nemopilema* by aligning the homeodomain sequences of human and fruit fly to the identified *Nemopilema* homeodomains. We considered only domains that were aligned to both the human and fruit fly. We also used this process for *Acropora*, *Hydra*, and *Nematostella* for comparison. Additionally, we added one Hox gene for *Acropora* and two Hox genes for *Hydra*, which are absent in NCBI gene set, though they were present in previous study [21, 59]. Hox and ParaHox genes of *Clytia hemisphaerica*, a hydrozoan species with a medusa stage, were also added based on a previous study [60]. Finally, a multiple sequence alignment of these domains was conducted using MUSCLE, and a FastTree [61] maximum-likelihood phylogeny was generated using the Jones–Taylor–Thornton (JTT) model with gamma option.

### *Wnt* gene subfamily analyses

*Wnt* genes of *Nematostella* and *Hydra* were downloaded from previous studies [23, 62], and those of *Acropora* were downloaded from the NCBI database. *Wnt* genes in *Nemopilema* were identified using the Pfam database by searching for “wnt family” domains. A multiple sequence alignment of *Wnt* genes was conducted using MUSCLE, and aligned sequences were trimmed using the trimAl program [63] with “gappyout” option. A phylogenetic tree was generated using RAxML with the PROTGAMMAJTT model and 100 bootstraps.

## Abbreviations

SMRT: Single molecule real time sequencing
TSLR: TruSeq synthetic long reads

## Declarations

### Availability of data and materials

The jellyfish genome project has been deposited at DDBJ/ENA/GenBank under the accession PEDN00000000. The version described in this paper is version PEDN01000000. Raw DNA and RNA sequence reads for *Nemopilema nomurai* and *Sanderia malayensis* have been submitted to the NCBI Sequence Read Archive database (SRA627560). All other data can be obtained from the authors upon reasonable request.

## Authors’ contributions

JB and SY supervised the project. YSC, JB, and SY planned and coordinated the project. HMK, JAW, YSC, SY, and JB wrote the manuscript. NayoungL, NayunL, YJJ, SW, KS, JCR, HSY, JHL, and SY prepared the samples, performed the experiments, and provided toxinological considerations. VL, AK, and MWK performed the gene evolutionary age analysis. HMK, SGP, YSC, YB, YJ, SJ, OC, JSE, and AM performed in-depth bioinformatics data analyses. All authors reviewed the manuscript and discussed the work.

## Ethics approval

This is not applicable.

## Acknowledgements

Korea Institute of Science and Technology Information (KISTI) provided us with Korea Research Environment Open NETwork (KREONET), which is the internet connection service for efficient information and data transfer.

## Additional files

**Additional file 1**: Supplementary figures, tables, and methods. This document contains additional supporting evidence for this study that are presented in form of supplemental figures and tables.

**Additional file 2**: Supplementary data. Protein domain annotation statistics.

**Additional file 3**: Supplementary data. List of gene clusters evolving faster in the *Nemopilema nomurai* genome.

**Additional file 4**: Supplementary data. Gene ontology and KEGG enrichment result of top 100 and 500 GC3 genes in *Nemopilema nomurai*.

**Additional file 5**: Supplementary data. Gene ontology enrichment result of highly expressed genes in *Nemopilema nomurai* and *Sanderia malayensis*.

